# Large-scale simulations reveal evolutionary constraints on intrinsically disordered regions imposed by full-length protein architecture

**DOI:** 10.64898/2026.02.27.708199

**Authors:** Yang Jiang, Xiaoyang Liu, Li Zhao, Zhong-Yuan Lu

**Author notes:** **Corresponding Authors:** Li Zhao, Zhong-Yuan Lu.

## Abstract

Intrinsically disordered regions (IDRs) are pervasive in eukaryotic proteomes and play central roles in regulatory complexity and evolutionary innovation. While their conformational ensembles have been extensively characterized in isolation, most long IDRs function within multidomain proteins, where domain architecture may impose structural and evolutionary constraints. Here, we perform proteome-scale molecular dynamics simulations of 14,283 human proteins containing IDRs, integrating the Calvados coarse-grained force field with an elastic network representation of structured domains to model full-length architectures. Focusing on 8,988 long IDRs (≥100 residues), we systematically compare conformational properties in isolated versus full-length contexts. We find that 2,733 IDRs (over 30%) undergo significant conformational shifts when embedded within their native protein architecture, revealing pervasive structural coupling between ordered and disordered regions. Notably, IDRs positioned centrally within proteins are evolutionarily biased toward compact and rigid conformations, whereas those with strong charge clustering preferentially adopt extended and flexible states in full-length contexts. Functional enrichment analyses further demonstrate that compact-rigid IDRs are overrepresented in DNA-binding proteins, while extended-flexible IDRs are enriched in RNA-binding functions, suggesting coordinated structural-functional specialization. Together, these findings support a model in which long IDRs do not evolve as independent polymeric segments, but instead co-evolve with structured domains as integrated architectural modules. Our results provide quantitative evidence that full-length protein organization imposes systematic conformational constraints on IDRs, revealing a previously underappreciated dimension of protein structural evolution and offering a new framework for understanding domain-disorder co-evolution across complex proteomes.

## Introduction

The human proteome comprises a vast array of intrinsically disordered regions. Many of these IDRs exceed 100 residues in length, alternate with structured domains, and perform complex biological functions that extend beyond simple linking or molecular recognition (Kato M, et al. 2012; van der Lee R, et al. 2014). Over the past two decades, certain prototypical long IDRs have attracted extensive attention due to their critical biological roles or their associations with significant medical conditions, such as neurodegenerative diseases (Boehning M, et al. 2018; Cui Q, et al. 2023). IDRs exhibit high flexibility and complex conformational ensembles, and their structural features cannot be obtained using conventional methods employed for structured proteins. Computational biology approaches provide an opportunity for the systematic understanding of the structural characteristics of IDRs. Researchers have begun analyzing statistical features of IDR structures across the proteome to better understand their functional mechanisms, leading to the development of coarse-grained molecular dynamics force fields (such as Calvados force field) suitable for simulating IDR structures (Cao F, et al. 2024).Studies have demonstrated that AlphaFold’s pLDDT (predicted local distance difference test) confidence score is a reliable indicator for predicting protein disorder (Tunyasuvunakool K, et al. 2021). By simulating predicted IDR segments from the human proteome based on pLDDT values using the Calvados force field, researchers have successfully elucidated the structural compactness characteristics of IDRs and identified correlations between this compactness and functions such as DNA binding and nuclear localization (Tesei G, et al. 2024).

At present, relevant studies only take into account the conformational information of IDR segments within proteins that is obtained through computational biology methods. Notably, in many well-studied proteins possessing such functions and localization signals, IDRs are frequently situated internally within long polypeptide chains containing multiple structured domains (Murthy AC, et al. 2021; Cui Q, et al. 2023). In addition, it has been confirmed that there is a cooperative relationship between structured domains and IDRs during the functional execution of biomolecular condensates (Strom AR, et al. 2024). These observations prompt further inquiry: within the human proteome, how do the conformations of long IDRs—regions characterized by high structural plasticity—become influenced by other structured domains of the same protein?

To address this question, we employed the Calvados force field combined with an elastic network approach to perform molecular dynamics simulations on 14,283 full-length human proteins containing IDRs (≥30 residues), alongside parallel simulations of the corresponding isolated IDR segments as controls. We focused on 8,988 long IDRs (>100 residues), comparing their conformational changes along the dimensions of compaction-extension and rigidity-flexibility under both simulation conditions. From the perspectives of sequence position and residue-level features, we identified the primary factors contributing to significant conformational influence from structured domains. We also analyzed the subcellular localization and molecular function distributions of proteins harboring IDRs with distinct types of conformational responses. This work adds valuable conformational data derived from dynamic simulations of IDRs in their full-length context and proposes a novel classification dimension for protein structural features based on the degree to which long IDRs are influenced by other parts of their host proteins.

## Results

### Full-length protein simulations and IDR structure characterization

Based on structural predictions from the AlphaFold database for human proteome, we identified the proteins containing intrinsically disordered regions (Jumper J, et al. 2021). A region was defined as an IDR if at least 30 consecutive residues exhibited pLDDT scores below 70. We identified 14,283 proteins containing IDRs, encompassing 29,150 IDR segments in total.

Structured domains were defined according to the CATH database (Sillitoe I, et al. 2021), utilizing the shortest domain length observed at the 99.9% confidence level (30 residues) as the threshold (Fig. 1A and Fig. 1B). Regions with at least 30 consecutive residues having pLDDT ≥70 were treated as structured, and elastic networks were applied based on their AlphaFold-predicted structures, thereby enabling full-length molecular dynamics simulations for all selected proteins.

**Figure 1.**
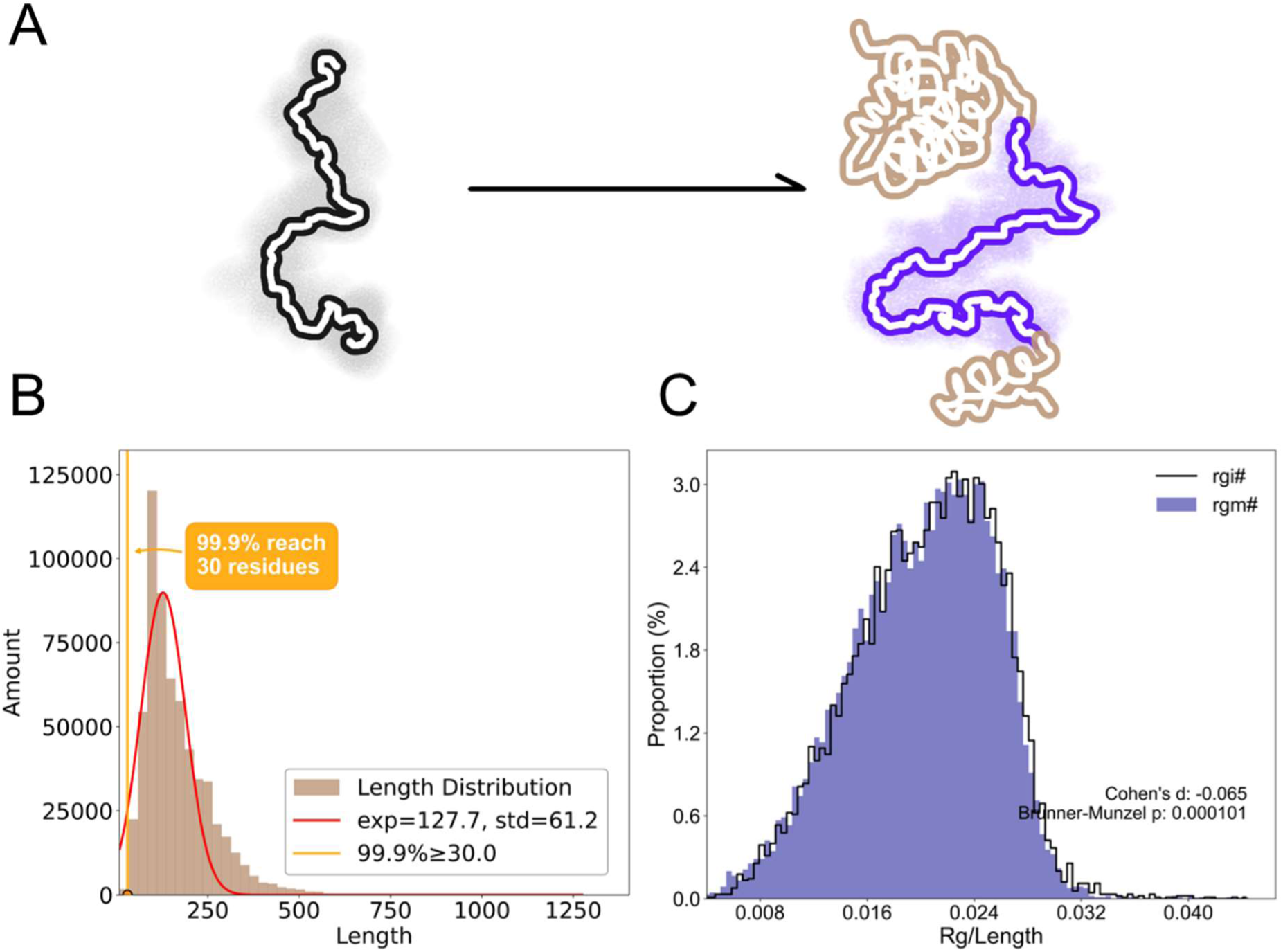
Schematic diagrams of the isolated IDR and full-length protein with elastic network added to the well-structured domains (A). Determination of the minimum sequence length of 30 residues for elastic network addition based on the CATH structural domain database (B). Conformational overall compression of IDRs with sequence lengths ranging from 100 to 1600, influenced by other domains in the full-length protein (C), where rgm# is Rg/length in full-length protein simulations, and rgi# is Rg/length in isolated IDR simulations.

Subsequently, we performed coarse-grained molecular dynamics simulations using the Calvados force field on both the full-length proteins and, in parallel, on their isolated IDR segments. To quantify conformational features, we utilized the ratio of the radius of gyration to sequence length (Rg/length). This parameter has been validated: predictions based on the Calvados force field align with experimental data for known representative proteins. Specifically, the mean Rg/length value derived from simulation trajectories characterizes the “compact-extended” dimension of IDR conformation, while its standard deviation reflects conformational fluctuations along the “rigid-flexible” dimension.

As shown in Fig. 1C, we focused our analysis on 8,988 long IDRs ranging from 100 to 1,600 residues. Analysis revealed that, within full-length proteins, these IDRs adopt more compact conformations compared to when simulated in isolation. This trend was different from all 29,150 IDR segments (SI Fig. S1).

### Classification of IDRs significantly influenced by full-length context into six categories across two dimensions

Numerous experimental studies have demonstrated that structured domains can significantly modulate IDR function (Xie W, et al. 2019; Loughlin FE, et al. 2019). To understand this phenomenon from a conformational perspective, we compared IDR conformations in full-length versus isolated simulations to enrich our understanding of human IDR structural traits. Due to the inherent flexibility of IDRs, the mean and standard deviation of Rg/length after equilibration still exhibit broad variation. To assess the impact of other protein regions, we calculated ΔRg/length_ave (change in compaction-extension) and ΔRg/length_std (change in rigidity-flexibility) between the two simulation setups. To determine significance, we employed the 95% confidence interval of inter-simulation differences observed in fully disordered proteins as the threshold.

As anticipated, over half of the 8,988 long IDRs showed no significant conformational difference between full-length and isolated simulations. Nevertheless, we identified 2,733 IDRs whose conformations were significantly influenced by the remainder of the protein. Two-dimensional analysis revealed that although most IDRs became more compact in the full-length context (Fig. 1C), 573 IDRs displayed a clear tendency toward extension, as shown in Fig. 2A.

**Figure 2.**
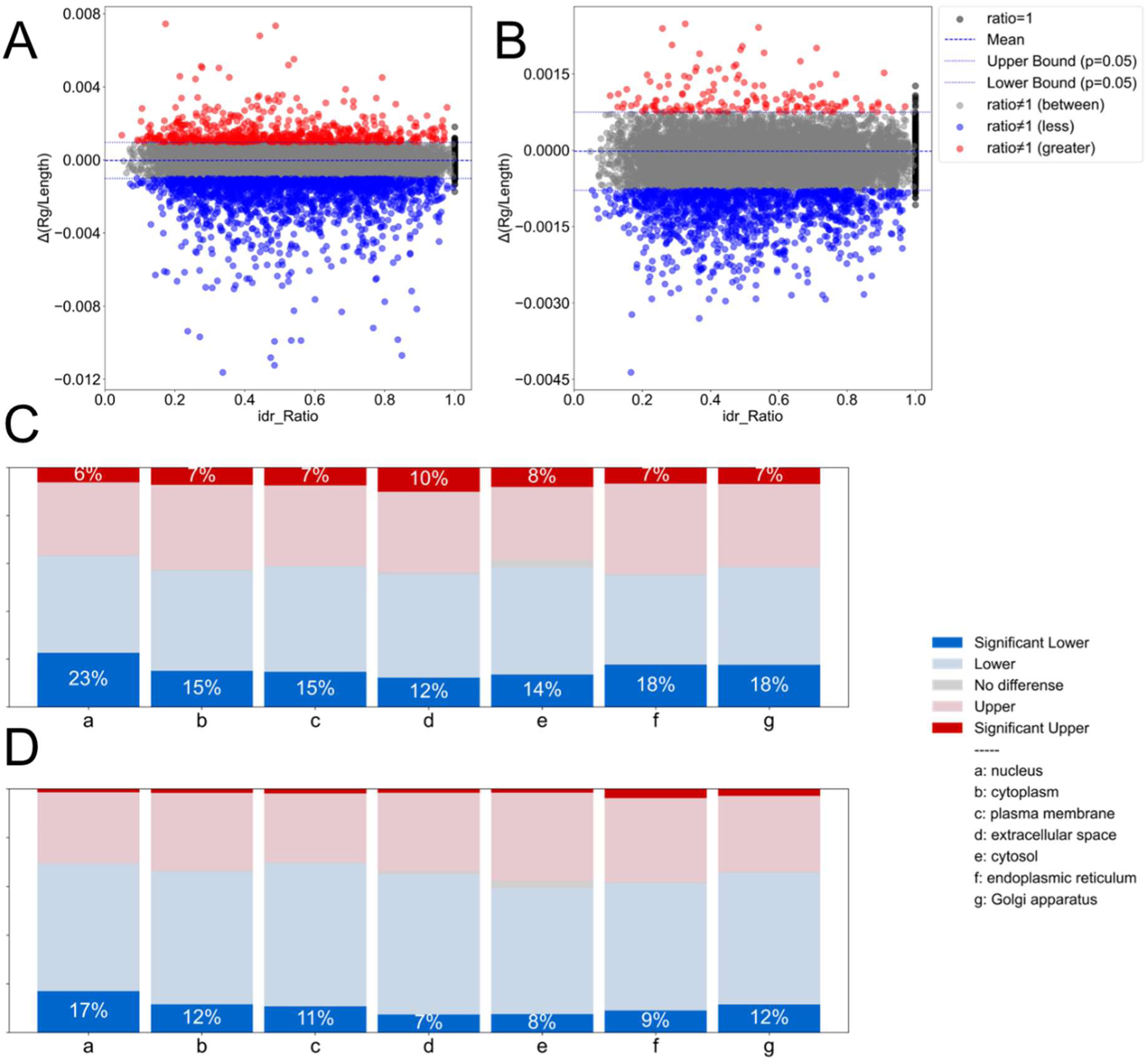
Comparison of conformational changes of the same long-chain IDRs (100-1600 residues) between full-length protein simulations and isolated IDR simulations from two dimensions. The average difference in Rg/length between full-length and isolated IDR simulations reflects compaction-extension dimensional changes, revealing 573 extended IDRs (red) and 1664 compacted IDRs (blue), in addition to those with negligible changes (gray) (A). The standard deviation difference in Rg/length reflects rigidity-flexibility dimensional changes, revealing 144 more flexible IDRs (red) and 1275 more rigid IDRs (blue), aside from those with negligible changes (gray) (B). In protein groups annotated with different subcellular localizations, IDRs influenced by other parts within full-length proteins exhibit varying compaction-extension dimensional effects distributed across groups, where light red represents IDRs in the gray region above the zero line in (A), and light blue represents IDRs in the gray region below the zero line in (A) (C). In protein groups annotated with different subcellular localizations, IDRs influenced by other parts within full-length proteins exhibit varying rigidity-flexibility dimensional effects distributed across groups, where light red represents IDRs in the gray region above the zero line in (A), and light blue represents IDRs in the gray region below the zero line in (A) (D).

Using ΔRg/length_ave, we categorized IDRs: those above the upper bound of the 95% confidence interval were labeled “more extended”, and those below the lower bound were labeled “more compact”. Gene ontology-based subcellular localization analysis demonstrated that both groups are distributed across all major cellular compartments in Fig. 2C, underscoring the importance of considering the full-length context when studying IDR function (Moreno P, et al. 2022). Similarly, based on ΔRg/length_std, IDRs above the upper bound were classified as “more flexible,” and those below as “more rigid” (Fig. 2B). The spatial distribution pattern of affected IDRs in this dimension closely mirrored that of the first, as shown in Fig. 2D.

Notably, within nuclear-localized IDRs, the proportions of “more compact” and “more rigid” categories were significantly higher than in other compartments (Fig. 2C and Fig. 2D). This aligns with previous reports indicating generally tighter conformations of nuclear IDRs (Tesei G, et al. 2024), suggesting that some nuclear proteins may utilize structured domains to further compress IDR conformations and restrict flexibility, thereby adapting to the densely packed molecular environment inside the nucleus.

By combining both ΔRg/length_ave and ΔRg/length_std, we classified significantly affected long IDRs into six distinct conformational types. A significant positive correlation was observed between the two dimensions (Fig. 3A): effects leading to compaction in full-length proteins typically coincided with reduced flexibility, whereas those promoting extension usually increased flexibility.

**Figure 3.**
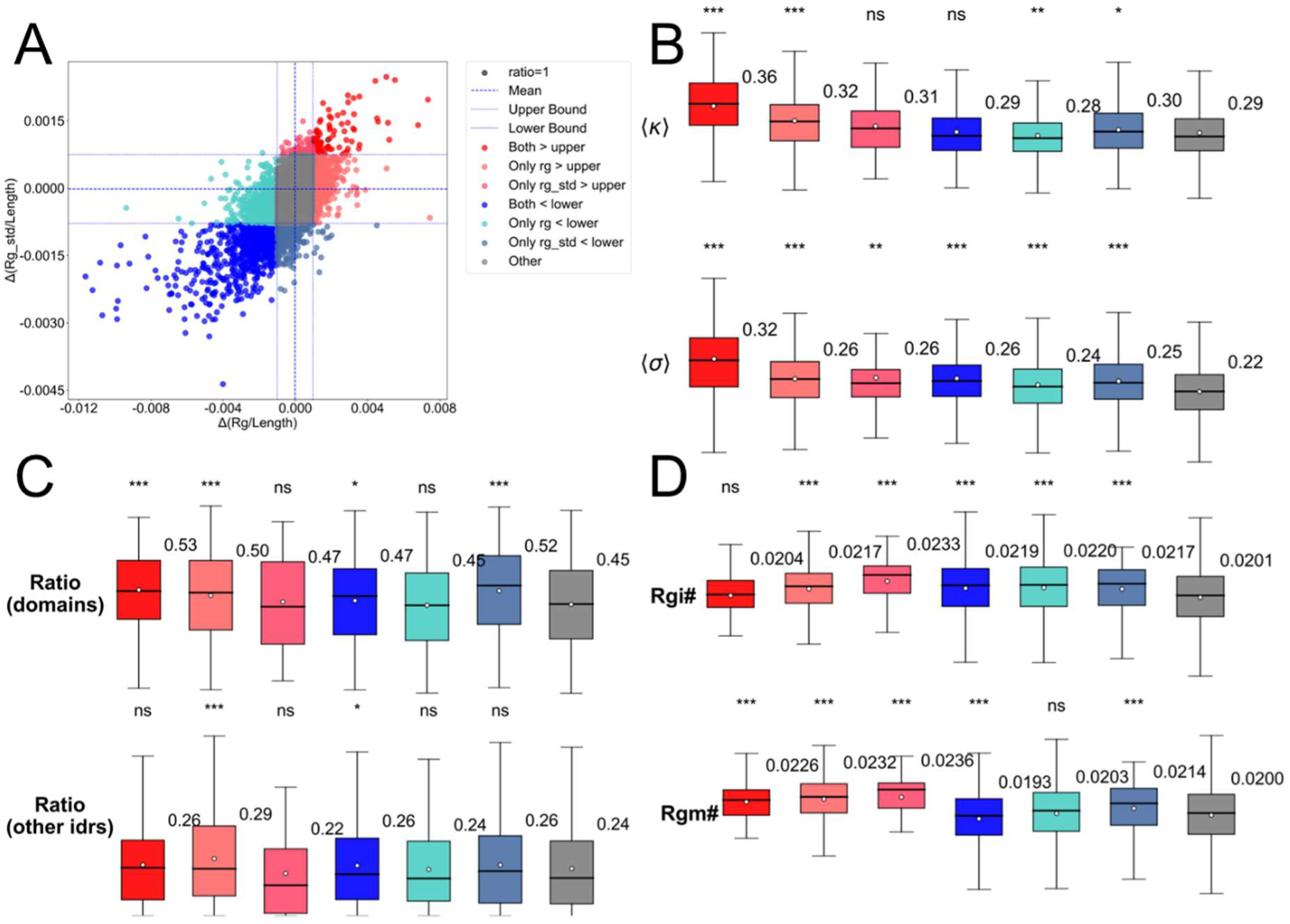
IDRs are classified into an insignificant-effect group and six significant-effect groups based on their response to other sequences in the full-length protein across two dimensions (A). The degree of charged residue clustering <κ> and the charged residue content <σ> for the insignificant-effect group and the six significant-effect groups (B). The proportion of structured domains and other IDRs in the full-length protein for IDRs belonging to the insignificant-effect group and the six significant-effect groups (C). The Rg/length in full-length protein simulations (rgm#) and in isolated IDR simulations (rgi#) for IDRs in the insignificant-effect group and the six significant-effect groups (D).

Moreover, the influence from other parts of the full-length protein could substantially alter absolute conformational compactness. Among the six classes, IDRs in the “more extended and more flexible” group were notably more expanded than other groups, while some “more rigid” IDRs also exhibited significant compaction (Fig. 3D). This confirms that, for many long IDRs, accurate characterization requires consideration of their conformation within the full-length protein system.

### Relationship between sequence features and conformational influence from structured domains

The clustering of charged residues within IDR sequences, particularly their non-uniform distribution, has been shown in multiple studies to drive chain compaction (Pesce F, et al. 2024; Tesei G, et al. 2024). Our results indicate that for specific types of long IDRs—particularly those becoming more extended—a higher degree of charge clustering correlates with a more extended conformation in the full-length context (Fig. 3B, SI Fig. S2). This implies that the influence of charged residues cannot be fully captured by simulating isolated IDR fragments alone; other components in the full-length protein may be essential for the complete functional expression of such IDRs. Further analysis revealed that, in addition to higher charge clustering, IDRs in the “more extended and more flexible” group tend to reside in proteins where structured domains occupy a larger fraction of the sequence (Fig. 3C). This suggests that the presence of structured domains and intrinsic sequence charge properties jointly shape the extended, flexible conformations of these IDRs.

Conversely, another abundant class of IDRs exhibited more compact and rigid conformations in the full-length context. These IDRs typically possess a higher overall proportion of charged residues but lack pronounced charge clustering. Notably, they are frequently located in the central regions of protein sequences—rather than near the N- or C-termini—a positional feature not observed in the more extended categories (Fig. 4A). This indicates that being positioned between multiple structured domains (i.e., centrally) may specifically promote a more compact and rigid conformation for such IDRs.

**Figure 4.**
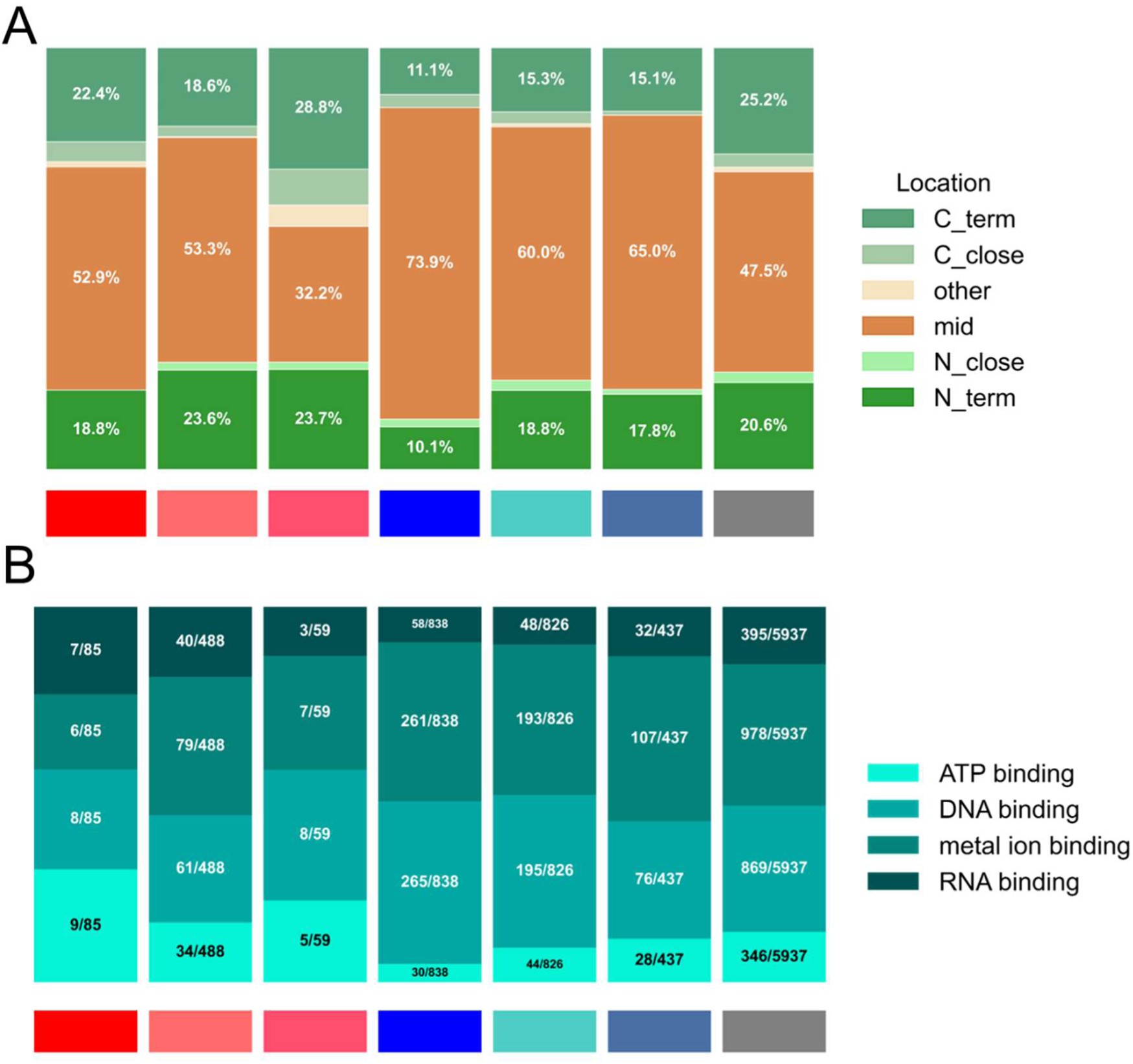
Cellular localization and molecular function distributions of IDRs in the insignificant-effect group and the six significant-effect groups were obtained through GO analysis. The main subcellular localizations are summarized as shown in panel (A), the color scheme corresponding to the insignificant-effect group and the six significant-effect groups can be found in Figure 3A. IDRs in both the insignificant-effect group and the six significant-effect groups are distributed across the four main subcellular localizations, with a notably higher proportion of more compact and more rigid IDRs localized in the nucleus compared to other categories (B).

### Cellular localization and function of IDRs significantly influenced by full-length context

We have demonstrated that IDRs significantly influenced by structured domains are widely distributed across major cellular compartments. To explore their biological implications, we compared gene ontology (GO) annotations of proteins containing unaffected IDRs versus those harboring one of the six significantly affected types.

Overall, the “more compact and more rigid” category showed strong enrichment for nuclear localization. Yet, nuclear-localized proteins were also abundant in other categories (Fig. 5A and Fig. 5B). Subcellular analysis revealed that all six types of significantly affected IDRs exist in both nuclear compartments (e.g., chromatin, nucleolus, nuclear bodies) and cytoplasmic organelles (e.g., endoplasmic reticulum, Golgi apparatus, mitochondria) (SI Fig. S5B).

**Figure 5.**
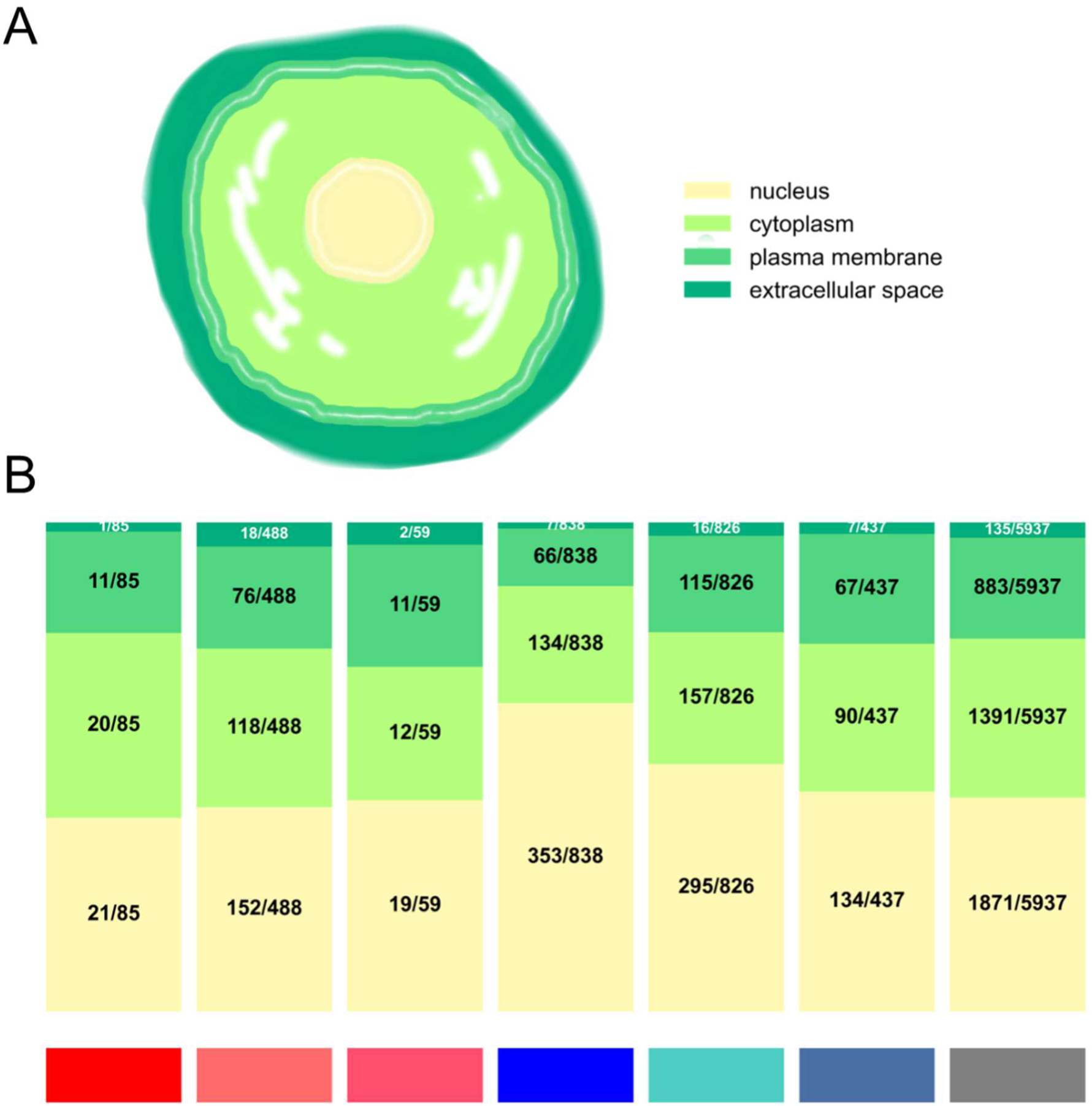
The proportion of IDRs in the insignificant-effect group and the six significant-effect groups varies across different positions within the full-length sequence (N-terminal, near N-terminal, middle, near C-terminal, C-terminal). Notably, the majority of IDRs that become more compact and more rigid are located in the middle of the full-length sequence (A). The proportions of IDRs in the insignificant-effect group and the six significant-effect groups involved in different binding functions vary. IDRs that are more compact and more rigid show a higher proportion annotated with DNA binding, while IDRs that are more extended and more flexible exhibit a higher proportion annotated with RNA binding (B).

Proteins in different categories exhibit distinct molecular binding functions. Specifically, IDRs in the “more compact and more rigid” group are enriched in proteins with DNA-binding activity, whereas those in the “more extended and more flexible” group are more commonly associated with RNA-binding functions (Fig. 4B). This suggests that structured domains involved in different nucleic acid interactions exert distinct influences on adjacent IDRs, implying that IDRs may co-evolve with structured domains to meet the overall functional requirements of the protein.

Furthermore, root-node tracing of unique GO terms in each category revealed that all six classes include unique components of nuclear protein complexes and enzyme activators. Particularly noteworthy is the discovery of several unique microtubule-organizing complex-related proteins exclusively within the “more extended and more flexible” IDR category (SI Fig. 6).

## Discussion

Intrinsically disordered regions play crucial roles in a range of key physiological processes. However, traditional experimental techniques struggle to resolve their dynamic structures and link them to specific functions, rendering computational simulations highly valuable for advancing our understanding. Thanks to prior work, coarse-grained modeling approaches for predicting and statistically analyzing IDR conformations have become established and accepted, providing initial insights into IDR structural statistics across the human proteome. Yet, in physiological process, for example, stress granules (Wheeler JR, et al. 2016) IDRs—especially long ones—typically function through complex interactions with other domains or IDRs within the same protein, a mechanism not yet fully understood. This study presents the first large-scale molecular dynamics effort to obtain conformational statistics of IDRs within full-length human proteins and systematically compares them with isolated IDR simulations. It thus effectively supplements IDR conformational data under conditions approximating physiological reality. We not only identify IDRs whose conformations are largely independent of the rest of the protein but, more importantly, classify six distinct types of long IDRs whose conformations are significantly shaped in different ways by other parts of their host proteins. These classifications offer new clues and analytical dimensions for future experimental studies on the structure and function of specific long-IDR-containing proteins.

The conformations of long IDRs are profoundly shaped by their protein environments—they may become more compact and rigid, or more extended and flexible. Our study reveals that sequence position contributes to the former, while charge clustering drives the latter. Notably, we observe that the direction of charge clustering effects on IDR conformation may reverse between isolated and full-length contexts, underscoring the necessity of studying IDRs in physiologically relevant settings. Moreover, all six classes of significantly affected IDRs are broadly distributed across cellular compartments, each exhibiting unique localization patterns. Their distinct preferences for different nucleic acid binding functions provide new insights into the functional coordination—and potential co-evolution—between structured domains and long disordered regions.

## Methods

### Structured domain treatment and IDR selection

The AlphaFold2 Database provided predicted structural data for the human proteome, including initial conformations for full-length protein simulations and per-residue pLDDT scores. In the full-length simulation setup, continuous sequences of 30 or more residues with pLDDT >70 were defined as structured domains. Elastic networks were applied to these regions to represent them within the coarse-grained Calvados force field, which is adapted for IDR simulations. Remaining regions were treated as disordered (no elastic network added). This definition follows the CATH database, utilizing the shortest recorded domain length at 99.9% confidence (30 residues).

For parallel simulations of isolated IDR sequences, continuous regions with pLDDT ≤70 were extracted, retaining only those with lengths of at least 30 residues. Initial structures were still based on AlphaFold predictions.

### Elastic network setup

Using the elastic network tool provided in Calvados3, elastic networks were applied to defined structured domains with a distance cutoff of 0.9 nm and a spring constant (k) of 700 kJ/(mol·nm^2^).

### Molecular dynamics simulations

Molecular dynamics simulations were conducted in the NVT ensemble at 37 °C using the Langevin integrator with a time step of 10 fs and a friction coefficient of 0.001 ps^−1^. The Cα-based model was implemented using Calvados3 parameters and functional forms as previously described. Salt-screened electrostatic interactions were modeled assuming a dielectric constant of 74.2 and an ionic strength of 0.1 M. Simulation box side length was set to 0.38 × (number of residues − 1) + 4.0 nm. Default energy minimization parameters were used. Each chain was simulated for 220 ns, with the first 20 ns simulation trajectory discarded, resulting in 200 conformations per protein. The molecular dynamics simulations were carried out using IPAMD software (Liu XY, et al. 2025).

Mean and standard deviation of Rg/length were computed from 200 trajectory frames. Comparisons were made by extracting corresponding IDR segments from full-length simulation trajectories, aligned with sequences used in isolated IDR simulations. In sequence feature calculations, positively charged residues include arginine and lysine, while negatively charged residues include aspartic acid and glutamic acid. In κ (score for the concentration distribution of charged residues) calculations, uncharged sequence segments were assigned a value of 0.

### GO analysis

Human proteome gene annotation data were obtained from https://current.geneontology.org/annotations/goa_human.gaf.gz. Annotations were indexed using UniProt IDs of full-length proteins. Root-node tracing of unique GO terms was performed using the ontology file available at http://purl.obolibrary.org/obo/go/go-basic.obo.

## Acknowledgments

This work is supported by the National Natural Science Foundation of China (22473048 and 22133002). L.Z. acknowledges valuable discussion with Prof. Yong Wang at Zhejiang University.

